# Unravelling the roadblocks to 1,2-propanediol biosynthesis in select solventogenic *Clostridium* species

**DOI:** 10.1101/2024.02.22.581430

**Authors:** Eric Agyeman-Duah, Santosh Kumar, Victor C. Ujor

**Affiliations:** Department of Food Science, University of Wisconsin-Madison, Babcock Hall, 1605 Linden Drive, Madison 53706 WI USA

**Keywords:** 1,2-propanediol, acetol, butanol, methylglyoxal, hydroxyacetone, lactaldehyde

## Abstract

**Background:** The compound 1,2-propanediol is an important industrial bulk chemical that has proven particularly recalcitrant to bio-production. Solvent-producing *Clostridium* species represent promising candidates for engineering 1,2-propaediol production. Co-production of 1,2-popanediol and butanol has the potential to improve the economics of the acetone-butanol-ethanol (ABE) fermentation.

**Results:** In this study, the methylglyoxal synthase gene (*mgsA*) from *Clostridium beijerinckii* NCIMB 8052 was homologously expressed in this organism. Additionally, a separate strain of *Clostridium beijerinckii* NCIMB 8052 was engineered by cloning and expressing *mgsA* and methylglyoxal/glyoxal reductase (*mgR*) from *Clostridium pasteurianum* ATCC 6013 as a fused protein linked by polyglycine linker in the former. Both strains of *C. beijerinckii* NCIMB 8052 failed to produce 1,2-propaneol. Instead, traces of acetol—the precursor of 1,2-propanediol—were detected in cultures of both strains. When the recombinant strains were exposed to acetol, both strains exhibited ∼100% acetol-to-1,2-propanediol conversion efficiency. Conversely, methylglyoxal supplementation led to the production of traces of acetol but not lactaldehyde or 1,2-propanediol. When wildtype *C. beijerinckii* NCIMB 8052, *C. pasteurianum* ATCC 6013 and *Clostridium tyrobutyricum* ATCC 25755 were challenged with methylglyoxal, *C. beijerinckii* produced ∼0.1 g/L (*S*)-(+)-1,2-Propanediol, while *C. tyrobutyricum* produced traces of lactate. *C. pasteurianum* produced neither 1,2-propanediol nor lactate. The wild types of all three species above exhibited ∼100% acetol-to-1,2-propanediol conversion efficiency. The recombinant strain of *C. beijerinckii* expressing fused MgsA and MgR from *C. pasteurianum* ATCC 6013 showed enhanced growth and solvent production, producing as high as 88% more butanol on both glucose and lactose than the control strain and the recombinant strain of the same organism expressing the native MgsA.

**Conclusions:** Recombinant and native strains of *C. beijerinckii*, *C. pasteurianum* and *C. tyrobutyricum* studied in this work exhibit extremely poor capacity to catalyze the conversion of the intermediates of the methylglyoxal bypass to 1,2-propanediol. This is indicative of lack of appropriate enzymes to catalyze the reactions from methylglyoxal to acetol or lactaldehyde. Inability to detect methylglyoxal in the recombinant strains harboring *mgsA* (both homologous and heterologous)— whereas the strain expressing both *mgsA* and *mgR* from *C. pasteurianum*, under the same promoter (P*adc*) produced higher concentrations of butanol—suggests that *C. beijerinckii* might possess a regulatory mechanism that limits the activity of methylglyoxal-producing MgsA. The protein product of *mgR* from *C. pasteurianum* represents a promising metabolic engineering candidate towards increasing butanol production.

## Introduction

Bio-production of chemicals from renewable feedstocks is central to efforts aimed at decarbonizing the economy. Given the enormous variety of chemicals derived from finite petroleum and their numerous applications, bio-production of chemicals that readily lend themselves to multifarious reactions are particularly attractive towards decarbonization. This is because; such compounds find assorted applications across the economy by serving as precursors of several other compounds. Thus, they can replace multiple petroleum-derived chemicals. The compound 1,2-propanediol (1,2-PD), fits this criterion. Notably, 1,2-PD is used in the production of unsaturated polyester resins, as an additive in food processing, in making building materials, as a solvent in the pharmaceutical industry, to produce non-ionic detergents and cosmetics, as an ingredient in hydraulic fluids and as an antifreeze/de-icing agent [1-4]. As at 2020, the global 1,2-PD market was valued at $US 0.37 billion, and is on track to reach $US ∼0.40 billion by 2026 [4]. However, 1,2-PD is produced solely from petroleum feedstocks via a process (chemical hydration of fossil fuel-based propylene) that is capital-intensive, consumes large amounts of energy and produces waste streams that pose a formidable threat to the environment, in addition to the associated CO_2_ emissions [1,4,5]. Furthermore, chemical synthesis of 1,2-PD generates racemic mixtures of the two stereoisomers of 1,2-PDO (*R*-1,2-PDO and *S*-1,2-PDO). Racemic mixtures of 1,2-PD are less amenable to separation and organic synthesis of chiral pharmaceuticals [1,4,6]. Therefore, bio-production promises an eco-friendlier and separation-pliable alternative to chemical synthesis of 1,2-PD.

It is important to note however, that despite a lot of promise, 1,2-PD has proven particularly recalcitrant to bio-production at levels that would justify commercialization. Remarkably, despite extensive efforts over the years, 9.0 g/L is the highest concentration of 1,2-PD reported to date via biological production, which was achieved with wildtype *Clostridium thermosaccharolyticum* [7]. This is largely ascribable to a number of bottlenecks that plague the 1,2-PD pathway, referred to as the methylglyoxal bypass [**Fig. 1**]. Specifically, methylglyoxal, a central intermediate in the natural 1,2-PD pathway is extremely toxic [8-11]. As a result, the methylglyoxal bypass is tightly regulated. Consequently, the 1,2-PD pathway is less metabolically favored, being only activated under strict conditions of carbon overflow and in some organisms, phosphate limitation [12-14]. Accordingly, methylglyoxal synthase (MgsA; **Fig. 1**) is inhibited by phosphate in some bacteria [14]. Given the cellular importance of phosphate, physiological conditions that would normally favor growth and biosynthesis of other chemicals inhibit 1,2-PD production in such bacteria. Therefore, improving bio-production of 1,2-PD hinges on better understanding of the metabolic bottlenecks of the methylglyoxal bypass. Thus far, efforts to increase bio-production of 1,2-PD have led to the construction of an artificial 1,2-PD pathway [**Fig. 1**;15]. Nevertheless, this approach has yet to increase 1,2-PD titer and yield.

**Fig. 1.**
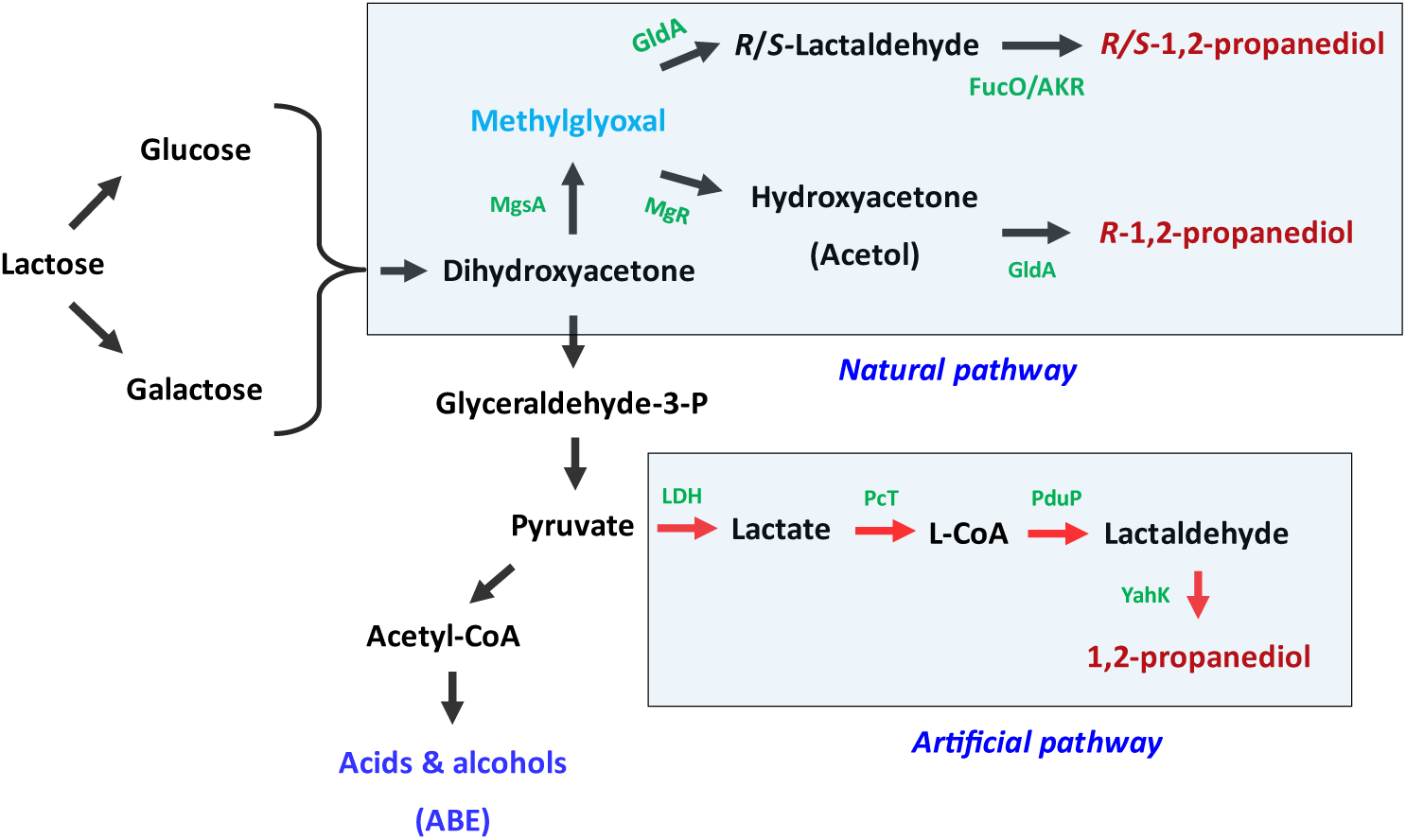
The 1,2-PD pathway (natural and artificially engineered). Lignocellulose-derived glucose and lactose from waste whey permeate may serve as feedstocks for 1,2-PD production. MgsA – methylglyoxal synthase, GldA – glycerol dehydrogenase, FucO – 1,2-PD oxidoreductase, AKR – aldehyde reductase, MgR – methylglyoxal reductase, LDH – lactate dehydrogenase, PcT - propionate CoA-transferase, PduP - aldehyde dehydrogenase, YahK - alcohol dehydrogenase.

In this study, we evaluated the potential of 1,2-PD production by the model butanol-producing bacterium, *Clostridium beijerinckii* NCIMB 8052 (hereafter referred to as *Cbei*), via the methylglyoxal bypass. The choice of *Cbei* was informed by a number of factors. First, *Cbei* utilizes a wide range of carbon sources including lignocellulose-derived glucose, xylose and arabinose, as well as lactose, the sugar content of whey permeate, an increasingly available feedstock for economical bio-production. Second, co-production of 1,2-PD and butanol—another chemical that has proven intractable to economical production—would improve the overall economics of butanol (via the acetone-butanol-ethanol – ABE) fermentation. Third, as with other solventogenic *Clostridium* species, *Cbei* has the inherent capacity to produce and tolerate solvents. Additionally, we investigated the bottlenecks to 1,2-PD production in *Clostridium pasteurianum* ATCC 6013 (hereafter, *Cpas*) and *Clostridium tyrobutyricum* ATCC 25755 (hereafter, *Ctyro*). The methylglyoxal bypass is a potential source of hydroxyacetone (acetol; **Fig. 1**), which is equally versatile as an industrial platform chemical. Acetol is used to synthesize propionaldehyde (acrolein), acetone, polyols and furan derivatives, as an aroma agent in food processing, in textile dyeing, and in cosmetics as a skin-tanning agent [16-17]. Cloning and expression of genes encoding enzymes of the 1,2-PD pathway in *Cbei* led to biosynthesis of acetol but not 1,2-PD. Subsequently, likely impediments to 1,2-PD biosynthesis in *Cbei*, *Cpas* and *Ctyro* were investigated. Conversely, co-expression of genes for glyoxal reductase (*mgR*: the protein product is putatively annotated to reduce glyoxal and methylglyoxal) and methylglyoxal synthase (*mgsA*) from *Cpas* in *Cbei* resulted 87% and 88% increases in butanol production on lactose and glucose, respectively, relative to the control strain or the strain of *Cbei* expressing the native *Cbei mgsA* only.

## Materials and methods

### Bacterial strains and culture media

The *Escherichia coli* strains Top10 and Dh5α were procured from New England Biolabs (Ipswich, MA, USA). *Cbei* and *Cpas* were procured from the American Type Culture Collection (Manassas, VA, USA) and maintained as spore suspensions in the laboratory as previously described [18]. *Ctyro* was kindly provided by Dr. Yi Wang (Auburn University, Auburn, AL USA). *Ctyro* was grown overnight in Tryptone-Glucose-Yeast extract (TGY) medium [18-19] and then stored as 500 μL aliquots in glycerol (30%, v/v) at -80 °C for future experiments.

### Construction of plasmids and strains

Primers used to generate plasmid constructs are presented in **Table 1**. All plasmid constructs were based on the pWUR459 [20]. Two strains of *Cbei* (*Cbei*_*mgsA* and *Cbei*_*mgsA*+*mgR*) were constructed. To construct *Cbei*_*mgsA*, the native methylglyoxal synthase gene (*mgsA*) in *Cbei* (Cbei_0026; 360 bp) was amplified using the primers mgsACb-F and mgsACb-R (**Table 1**), designed to introduce *Apa*I, *Xho*1 and ribosome binding (RBS) sites into the resulting amplicon. The PCR product was digested with *Apa*I and *Xho*I and then purified (Wizard® SV Gel, Promega, WI, USA). The eluted fragment was ligated downstream of an inducible acetoacetate decarboxylase promoter (P*adc*) from *Cbei* in pWUR459, digested similarly with *Apa*I and *Xho*I. The resulting plasmid construct designated as pWUR459-*mgsA* was chemically transformed into *E. coli* Dh5α and selected on antibiotic plates (ampicillin; 100 µg/ml). Colony PCR was performed to screen for positive transformants and recombinant plasmid was isolated and validated by sequencing (using M13 forward & reverse primers, Eurofins Genomics, Louisville, KY, USA). Plasmid constructs were purified and electroporated into *Cbei* in an anaerobic chamber (Coy Laboratory Products Inc., Grass Lake, MI, USA) according to a previously described protocol [21]. After transformation, the cells were grown in TGY broth in the anaerobic chamber for 8 h before transferring to antibiotic-containing TGY agar plates. The anaerobic chamber was maintained at 35 ± 1 °C. Transformants were transferred onto TGY agar (0.5%, w/v agar) antibiotic selection plates supplemented with erythromycin (25 µg/ml). Subsequently, three colonies were separately inoculated into TGY broth supplemented with erythromycin (25 µg/ml) and grown anaerobically for 6-8 days, which allowed the cells to sporulate. Afterwards, the spores were washed five times with sterile distilled water, collected and stored in 4 °C for future experiments.

**Table 1.**
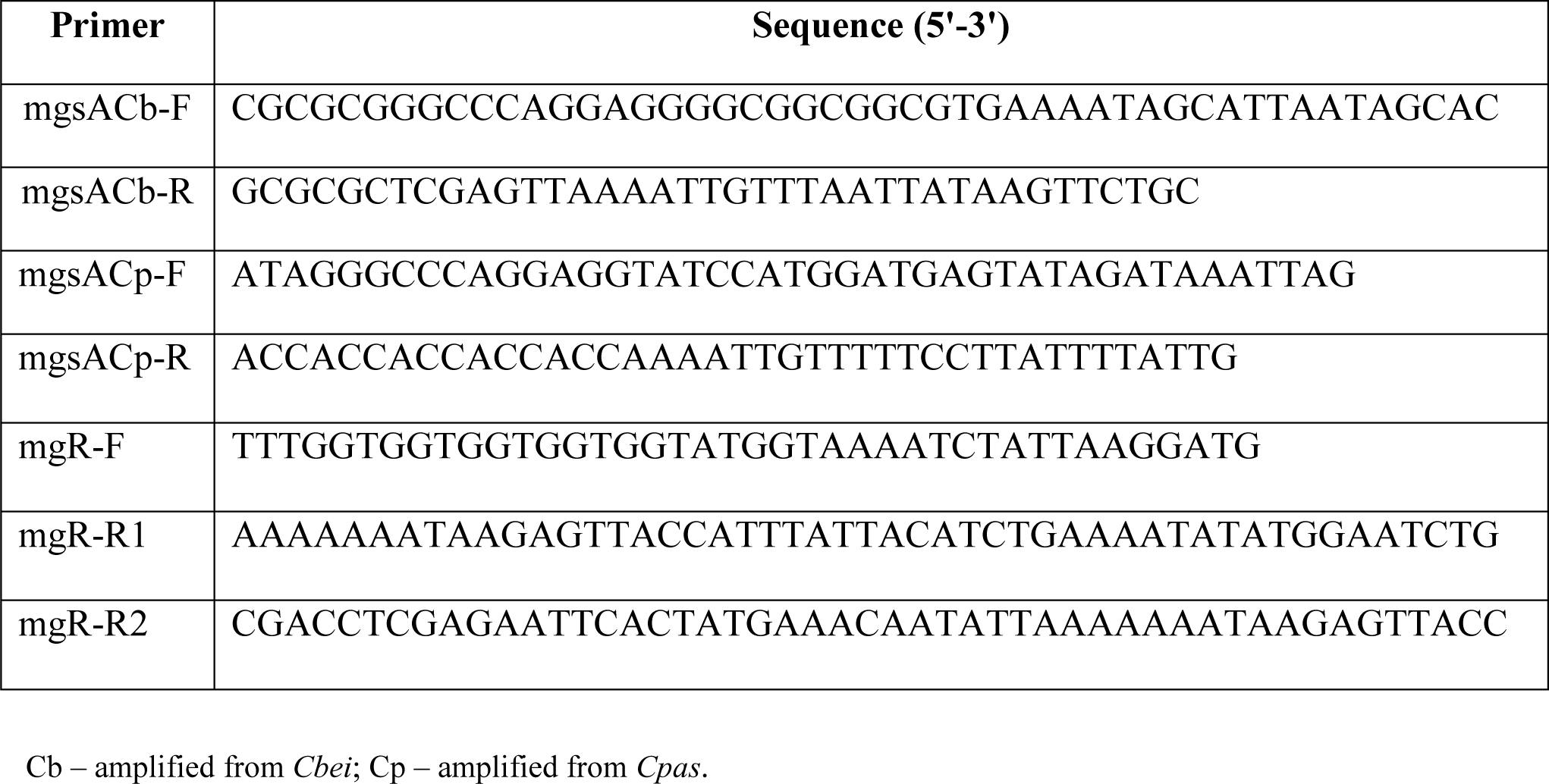
Primers used to construct *Cbei* clones bearing select genes of the 1,2-PD pathway

To construct *Cbei*_*mgsA+mgR*, *mgsA* and glyoxal reductase gene (*mgR*, annotated to reduce glyoxal and methylglyoxal) from *Cpas* were tethered together to encode a fusion protein including methylglyoxal synthase (MgsA) and glyoxal reductase (MgR), linked by a polyglycine linker. Because of the extreme toxicity of methylglyoxal, we rationalized that linking MgsA and MgR would enable more rapid detoxification of the catalytic product of MgsA (*i*.*e*., methylglyoxal). The gene, *mgsA* (AQ4_RS06990) from *Cpas* was amplified using the primers mgsACp-F and mgsACp-R (**Table 1**). The gene, *mgR* (AQ3_10695) was amplified with mgR-F and mgR-R1. Afterwards, *mgsA* and *mgR* were linked using splicing by overlap-extension (SOE) PCR with the primers mgsACp-F and mgsR-R2 (**Table 1**). The resulting amplicon was purified and ligated into pWUR459 downstream of P*adc* as earlier described. Afterwards, the construct was purified, validated by sequencing as described earlier and then, transformed into *E. coli* Top10. Subsequently, *Cbei* was transformed with the construct and colonies were picked and grown to spores as described earlier. Of the three colonies of *Cbei* picked from antibiotic agar plate, the colony with the most robust growth characteristic for each construct was selected for further studies.

To construct a control strain of *Cbei*, after plasmid sequencing, a gene lying downstream of the P*adc* promoter in pWUR459 was removed alongside P*adc* by restriction digestion with *Spe*I and *Xho*I. Afterwards, the linear plasmid was blunted by digestion with Quick Blunting™ Kit (NEB, USA) and ligated to re-circularize the plasmid. Re-circularized promoterless plasmid was designated as p459 and electroporated into *Cbei* as described above to obtain *Cbei*_p459.

### Cultivation of recombinant *Cbei* strains

Fermentations were conducted with *Cbei*_*mgsA*, *Cbei*_*mgsA*+*mgR* and *Cbei*_p459 in three different media including: (a) glucose (60 g/L), (b) glucose (60 g/L) supplemented with ZnSO_4_.7H_2_O (0.01 g/L) and (c) lactose (50 g/L). Because glycerol dehydrogenase (GldA), which catalyzes the conversion of methylglyoxal to lactaldehyde or acetol to 1,2-PD is zinc-dependent, ZnSO_4_.7H_2_O was supplemented to the glucose-based medium to determine if this might expedite methylglyoxal transformation to 1,2-PD. Each fermentation medium above was supplemented with P2 mineral, vitamin and buffer stocks as previously described [18]. Pre-cultures were prepared by growing the cultures in TGY as previously described [18]. Fermentation cultures of recombinant *Cbei* strains were supplemented with erythromycin 15 μg/ml. The fermentation media were inoculated with 6% (v/v) pre-culture. Samples (2 ml) were drawn every 12 h and analyzed for optical density, pH, 1,2-PD, acetol, acetone, butanol, ethanol, acetate and butyrate concentrations.

### Acetol and methylglyoxal challenge

To assess the rates of transformation and degrees of stress presented by the intermediates of 1,2-PD biosynthesis, fermentation in a glucose (60 g/L) medium was supplemented with acetol and methylglyoxal. *Cbei*_*mgsA*, *Cbei*_*mgsA*+*mgR* and *Cbei*_p459 were grown in the fermentation medium supplemented with erythromycin (15 μg/ml) for 12 h before methylglyoxal (0.5 g/L) addition. On the other hand, acetol (5 g/L) was supplemented at 0 h. Additionally, to determine whether the gridlock associated with 1,2-PD biosynthesis is limited to *Cbei*, glucose-grown cultures of wildtype *Cbei*, *Cpas* and *Cytro* were challenged with methylglyoxal (0.85 g/L added at 12 h) and acetol (0.2 and 5 g/L added at 0 h). The cultures were observed for growth and samples were drawn every 12 – 24 h (with the exception of samples taken 6 h after methylglyoxal supplementation) and assessed for methylglyoxal, 1,2-PD and acetol concentrations. Since wildtype *Cbei*, *Cpas* and *Ctyro* were not exposed to additional antibiotic stress, a higher methylglyoxal concentration (0.85 instead of 0.5 g/L) was added to cultures of the native strains.

### Analytical methods

Orion Star A214 pH meter (ThermoFisher Scientific, Waltham, MA, USA) was used to monitor culture pH during fermentation. Cell growth was determined as optical density (OD_600_ nm) using an Evolution 260 Bio UV/Visible spectrophotometer (ThermoFisher Scientific, Waltham, MA, USA). The concentrations of 1,2-PD, acetol, acetone, butanol, ethanol, acetate and butyrate were measured using a Shimadzu GC-2010 Plus gas chromatography (Shimadzu, Columbia, MD, USA) equipped with a fame ionization detector (FID), a Shimadzu AOC-20i auto injector, and a Shimadzu SH-PolarWax crossbond carbowax polyethylene glycol column [30 m (length), 0.25 mm (internal diameter), and 0.25 μm (film thickness)] as previously reported [18]. Additionally, gas chromatography-mass spectrometry (GC-MS) was used to qualitatively and in some cases, quantitatively screen culture samples for the presence of methylglyoxal, 1,2-PD, acetol and related metabolites. Sample preparation and GC-MS were conducted according to a previously reported method [18]. Glucose concentrations in the cultures were quantified using a YSI 2900 Series biochemistry analyzer (YSI, Yellow Springs, OH, USA) equipped with glucose/lactate calibrator, and configured with YSI 2357 buffer and the YSI 2365 glucose oxidase enzyme membrane according to manufacturer’s manual. Samples for glucose analysis were diluted 10x in 15% (w/v) saline and filtered through 0.22 µm sterile PTFE filter with 13 mm diameter (CELLTREAT, Pepperell, MA, USA). A total of 25 µL was aspirated for analysis.

Lactose concentrations were analyzed by HPLC. Samples of culture supernatant were filtered to remove cells and particulates as described for glucose analysis above. The samples were then diluted 1:10 before analysis in HPLC grade water (ThermoFisher Scientific, Waltham, MA, USA). The samples are then analyzed by an Agilent 1260 infinity HPLC system with a quaternary pump, chilled (4 °C) autosampler, vacuum degasser, and refractive index detector (Agilent Technologies, Inc., Palo Alto, CA). The HPLC column was an Aminex HPX-87H with Cation-H guard column 300×7.8mm (BioRad, Inc. Hercules, CA). The mobile phase was 0.02N H_2_SO_4_ with a flow rate of 0.500 mL/min. The column and detector were maintained 50 °C. Injection volume was 50 µl and the run time was 30 min. Instrument control, data collection and analysis and results calculation awere done using Chem Station V. B04.03 software (Agilent Technologies, Inc., Palo Alto, CA)

## Results

### Production of 1,2-PD and acetol

Cloning and expression of *mgsA* from *Cbei* and a fusion protein compromising the products of *Cpas mgsA* and *mgR* did not result in 1,2-PD production in *Cbei*. However, qualitative GC-MS analysis showed the presence of acetol—the precursor of 1,2-PD—in the culture broth extracts of the recombinant strains of *Cbei* under varying growth conditions (**Table 2**). Acetol was detected in cultures of *Cbei*_*mgsA* and *Cbei*_*mgsA*+*mgR* in all the media tested, with and without methylglyoxal and acetol supplementation, except when grown on glucose alone [without the addition of ZnSO_4_.7H_2_O (0.01 g/L), methylglyoxal or acetol]. Conversely, acetol was detected in the cultures of *Cbei*_p459 only when grown on glucose or lactose, without any culture supplements. Phosphate is an allosteric inhibitor of MgsA in some organisms [14,26]. Therefore, we grew the recombinant strains of *Cbei* in phosphate-limited media. However, cultivation of the recombinant strains of *Cbei* in phosphate-limited media did not lead to 1,2-PD production (data not shown). Interestingly, methylglyoxal challenge did not result in 1,2-PD production in all the recombinant strains of *Cbei* studied. Acetol challenge however, led to 1,2-PD production in all three recombinant strains of *Cbei*. In fact, careful examination of the results showed ∼100% conversion of acetol to 1,2-PD (data not shown). Further, growth on lactose led to acetol production in all three strains of *Cbei*.

**Table 2.** Detection of 1,2-PD and related metabolites in cultures of recombinant strains of *Cbei,* with and without methyl glyoxal and acetol supplementation.

| Strain | Methylglyoxal | Acetol | 1,2-PD |
| --- | --- | --- | --- |
| **Glucose** |  |  |  |
| *Cbei_p459* | - | + | - |
| *Cbei_mgsA* | - | - | - |
| *Cbei_mgsA* + mgR | - | - | - |
| **Glucose + 0.01 g/L ZnSO<sub>4</sub>·7H<sub>2</sub>O** |  |  |  |
| *Cbei_p459* | - | - | - |
| *Cbei_mgsA* | - | + | - |
| *Cbei_mgsA* + mgR | - | + | - |
| **Lactose** |  |  |  |
| *Cbei_p459* | - | + | - |
| *Cbei_mgsA* | - | + | - |
| *Cbei_mgsA* + mgR | - | + | - |
| **Glucose + 0.5 g/L methylglyoxal** |  |  |  |
| *Cbei_p459* | - | - | - |
| *Cbei_mgsA* | + | + | - |
| *Cbei_mgsA* + mgR | - | + | - |
| **Glucose + 5 g/L acetol** |  |  |  |
| *Cbei_p459* | - | - | + |
| *Cbei_mgsA* | - | + | + |
| *Cbei_mgsA* + mgR | - | + | + |

Samples were analyzed at 24 and 48 h

As shown in **Fig. 2**, *Cbei*_*mgsA*+*mgsR* and *Cbei*_p459 exhibited similar growth profiles with glucose and glucose supplemented with acetol (5.0 g/L). In both cases, when compared to *Cbei*_*mgsA*, both strains showed higher cell density, albeit only in the latter stages of fermentation (**Fig. 2A & C**). At the end of fermentation (108 h) in cultures grown on glucose alone, the optical densities (OD_600 nm_) of *Cbei*_*mgsA* reduced ∼18-fold (relative to the maximum OD achieved), whereas those of *Cbei*_p459 and _*mgsA*+*mgR* reduced 1.5- and 1.6-fold, respectively. In cultures grown on glucose supplemented with acetol, the reductions in OD_600 nm_ for all three strains were 1.7-, 3.4 and 19-fold for *Cbei*_ *mgsA*+*mgR*, _p459, and _*mgsA*, respectively, when compared to the maximum OD_600 nm_ (**Fig. 2C**) achieved by cultures of the respective strains. With methylglyoxal supplementation, at the end of fermentation, OD_600 nm_ reduced 2.41-, 3.4- and 27-fold, respectively, for *Cbei*_ *mgsA*+*mgR*, _p459, and _*mgsA* (**Fig. 2B**). However, earlier growth profiles (0 – 60 h) showed that *Cbei*_*mgsA* had similar growth patterns as *Cbei*_p459 and grew considerably better than *Cbei*_*mgsA*+*mgR* (34% higher maximum OD_600 nm_).

**Fig. 2.**
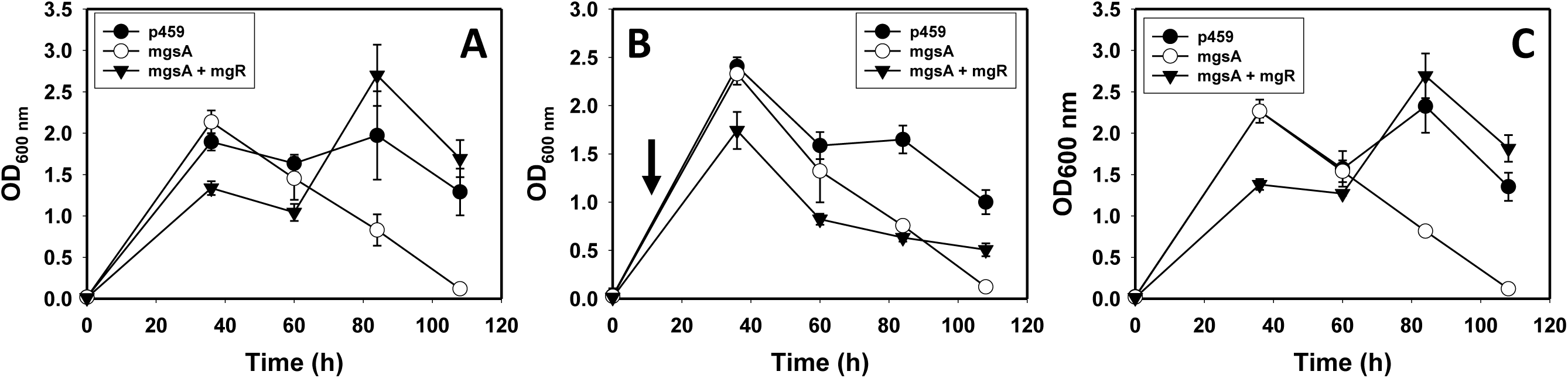
The growth profiles of *Cbei*_p459, _*mgsA* and _*mgsA* + *mgR* in glucose-based medium supplemented with methylglyoxal (0.5 g/L) and acetol (5 g/L) at 12 and 0 h, respectively, relative to un-supplemented cultures (glucose only control). A. glucose only cultures; B. glucose supplemented with methylglyoxal; C. glucose supplemented with acetol. Arrow (panel B) indicates addition of methylglyoxal at 12 h.

In an effort to establish whether *Cbei* is impaired for transforming the upstream intermediates of the methylglyoxal bypass (methylglyoxal and acetol) to 1,2-PD, we challenged wildtype *Cbei*, *Cpas* and *Ctyro* with methylglyoxal or acetol. At the concentrations tested (0.2 and 5.0 g/L supplemented at 0 h), acetol did not exert significant impediment to the growth of any of the three species studied (**Fig. 3**). Both 0.2 and 5.0 g/L acetol were efficiently transformed to 1,2-PD by all three species of *Clostridium* studied. While all the three species of *Clostridium* studied demonstrated efficient capability to transform acetol to 1,2-PD, *Ctyro* exhibited a slightly higher transformation rate than *Cbei* and *Cpas* (**Fig. 4**). After 24 h of acetol supplementation, the concentration of acetol decreased 92% in the cultures of *Ctyro* (**Fig 4A**). In parallel, the concentrations of acetol decreased 87% and 85% in the cultures of *Cbei* and *Cpas*, respectively, with a corresponding increase in 1,2-PD concentrations for all three species studied (**Fig 4B**).

**Fig. 3.**
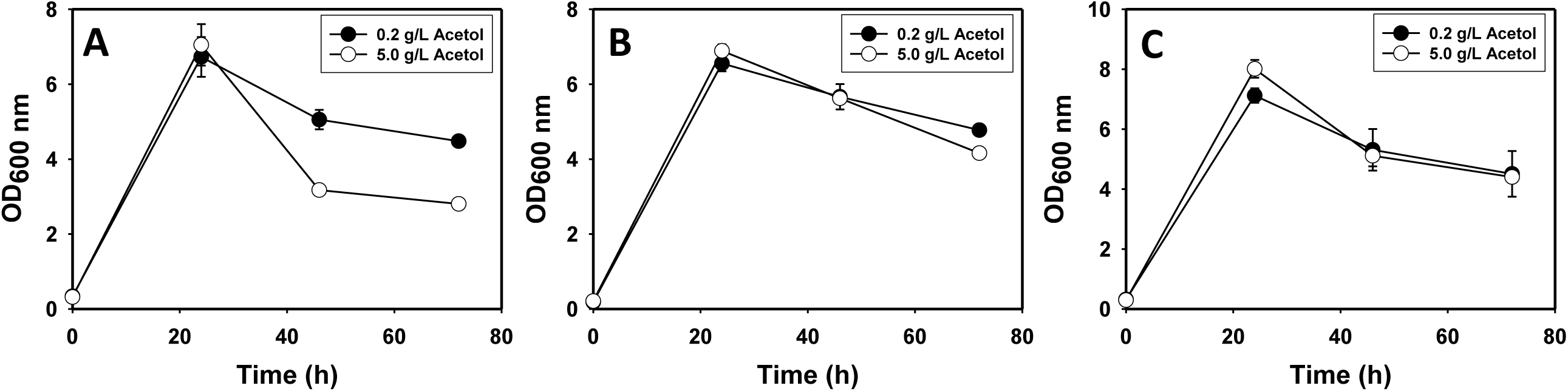
The growth profiles of *Cbei*, *Cpas* and *Ctyro* in glucose-based medium supplemented with acetol (5 g/L) at 0 h. A. *Cbei*; B. *Cpas*; C. *Ctyro*.

**Fig. 4.**
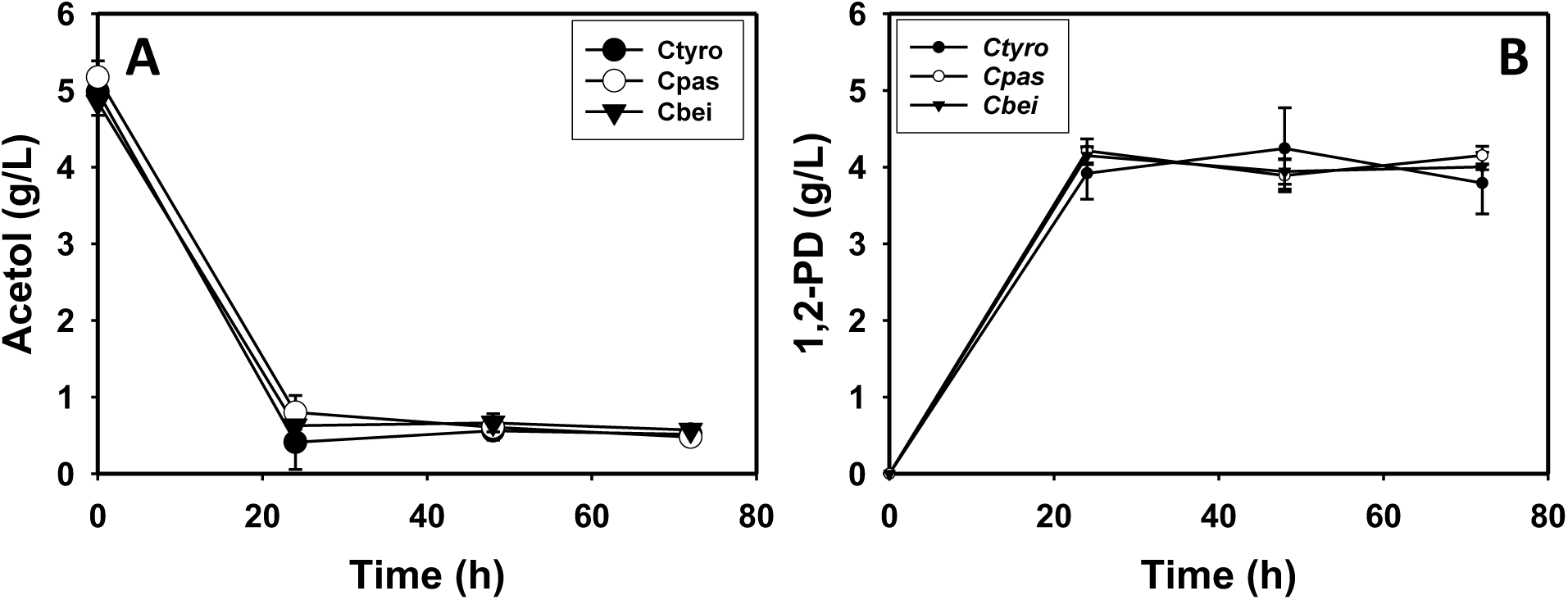
The concentrations of acetol and 1,2-PD in cultures of wildtype *Cbei*, *Cpas* and *Ctyro* supplemented with 5.0 g/L acetol at 0 h. A. Acetol concentrations; B. 1,2-PD concentrations.

When methylglyoxal (0.85 g/L) was added to cultures of *Cbei*, *Cpas* and *Ctyro*, drastic reduction in OD_600 nm_ was observed for all three species (**Fig. 5**). Specifically, 12 h after methylglyoxal addition, growth reduced ∼54%, ∼43% and 11% for *Cbei*, *Cpas* and *Ctyro*, respectively, when compared to the control cultures not supplemented with methylglyoxal. Among the three *Clostridium* species challenged with methylglyoxal, only *Cbei* produced 1,2-PD [specifically, (*S*)-(+)-1,2-PD; **Table 3**]. (*S*)-(+)-1,2-PD was detected by GC-MS and quantified by GC (0.08 g/L). Lactic acid was detected in the culture samples of *Ctyro*, whereas acetol was detected in the cultures of all three species tested. Residual methylglyoxal was detected only in the cultures of *Ctyro*.

**Fig. 5.**
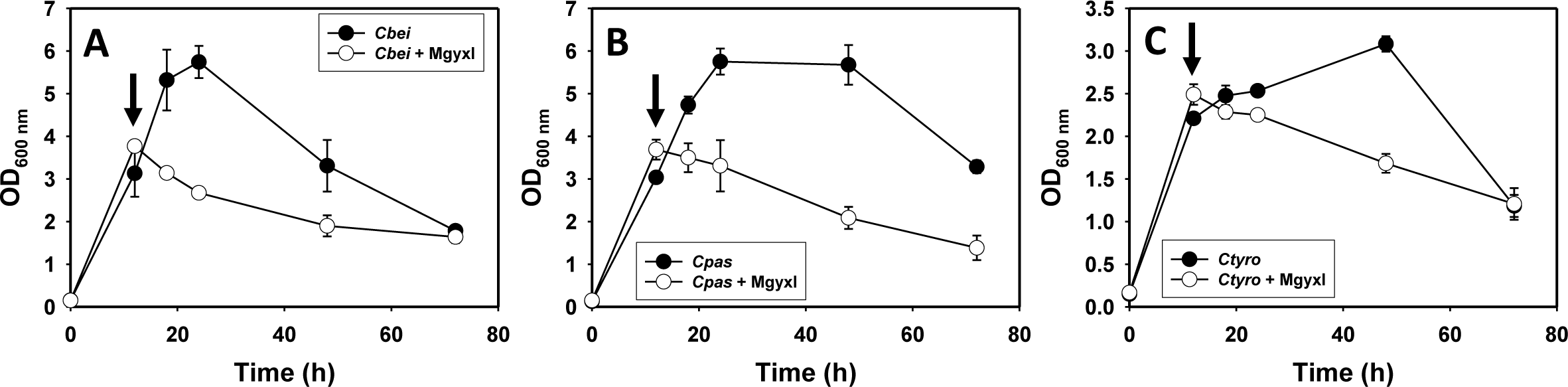
The growth profiles of *Cbei*, *Cpas* and *Ctyro* challenged with 0.85 g/L methylglyoxal (Mgyxl) at 12 h, relative to the un-supplemented cultures. A. *Cbei*, B. *Cpas*, C. *Ctyro*. Arrows indicate addition of methylglyoxal.

**Table 3.**
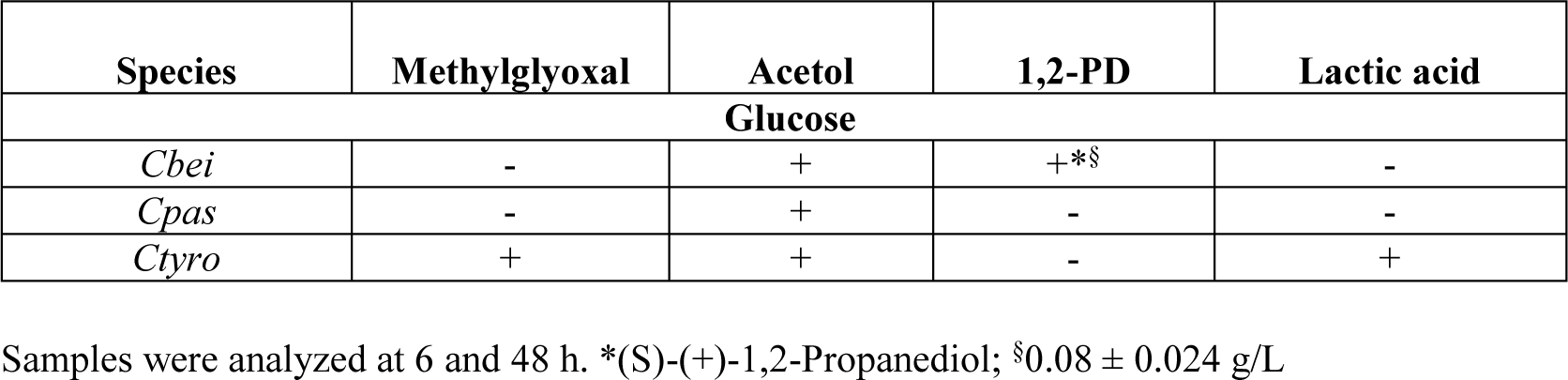
Detection of 1,2-PD and related metabolites in glucose-grown cultures of wildtype *Cbei*, *Cpas* and *Ctyro* supplemented with methylglyoxal.

| Species | Methylglyoxal | Acetol | 1,2-PD | Lactic acid |
| --- | --- | --- | --- | --- |
| Glucose |  |  |  |  |
| *Cbei* | - | + | +*§ | - |
| *Cpas* | - | + | - | - |
| *Ctyro* | + | + | - | + |

Samples were analyzed at 6 and 48 h. \*(S)-(+)-1,2-Propanediol; §0.08 ± 0.024 g/L

### Acetone-butanol-ethanol (ABE) production

Co-production of an appreciable concentration of 1,2-PD (or acetol) and butanol—the primary product of *Cbei*—would add value to ABE fermentation. Thus, *Cbei*_*mgsA*, *Cbei*_*mgsA*+*mgR* and *Cbei*_p459 were assessed for ABE production. Although *Cbei*_*mgsA*+*mgR* exhibited a slower growth rate on glucose, relative to *Cbei*_p459 and _*mgsA*, it eventually achieved a significantly higher maximum OD_600 nm_ (∼50% higher) than the other two strains (**Fig. 6A**). This translated to ∼73% and 48% higher butanol titer than *Cbei*_p459 and _*mgsA*, respectively (**Fig. 6B**). Accordingly, ABE titers in cultures of *Cbei*_*mgsA*+*mgR* were 72% and 52% greater than the titers detected in the cultures of *Cbei*_p459 and _*mgsA*, respectively (**Fig. 6C**; **Table 3**).

**Fig. 6.**
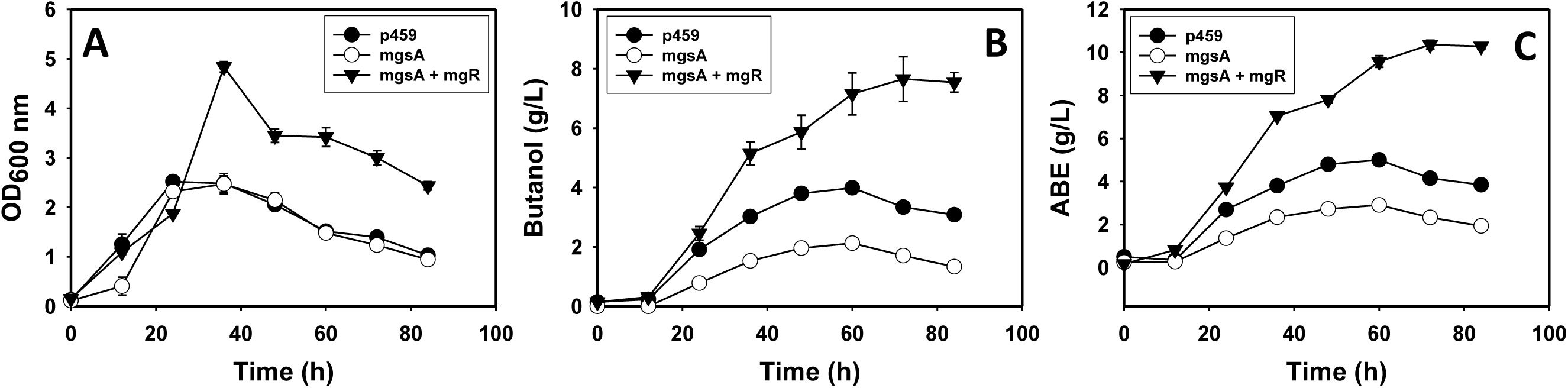
Growth, butanol and ABE profiles of recombinant strains of *Cbei* grown on glucose. A. OD_600 nm_; B. butanol concentrations; C. ABE concentrations.

The degree of acid (acetic and butyric acids) re-assimilation by solventogenic *Clostridium* species directly impacts butanol and to some degree, ABE production. Notably, the acetate and butyrate profiles of *Cbei*_*mgsA*+*mgR* suggest superior acid re-assimilation, relative to the other strains studied (**Fig. 7A**). Acetate concentrations in the cultures of *Cbei*_*mgsA*+*mgR* were at least ∼20% and 62% lower than those in the cultures of *Cbei*_p459 and _*mgsA*, respectively, whereas the butyrate concentrations were 48% and 84% lower, respectively (**Fig. 7B**).

**Fig. 7.**
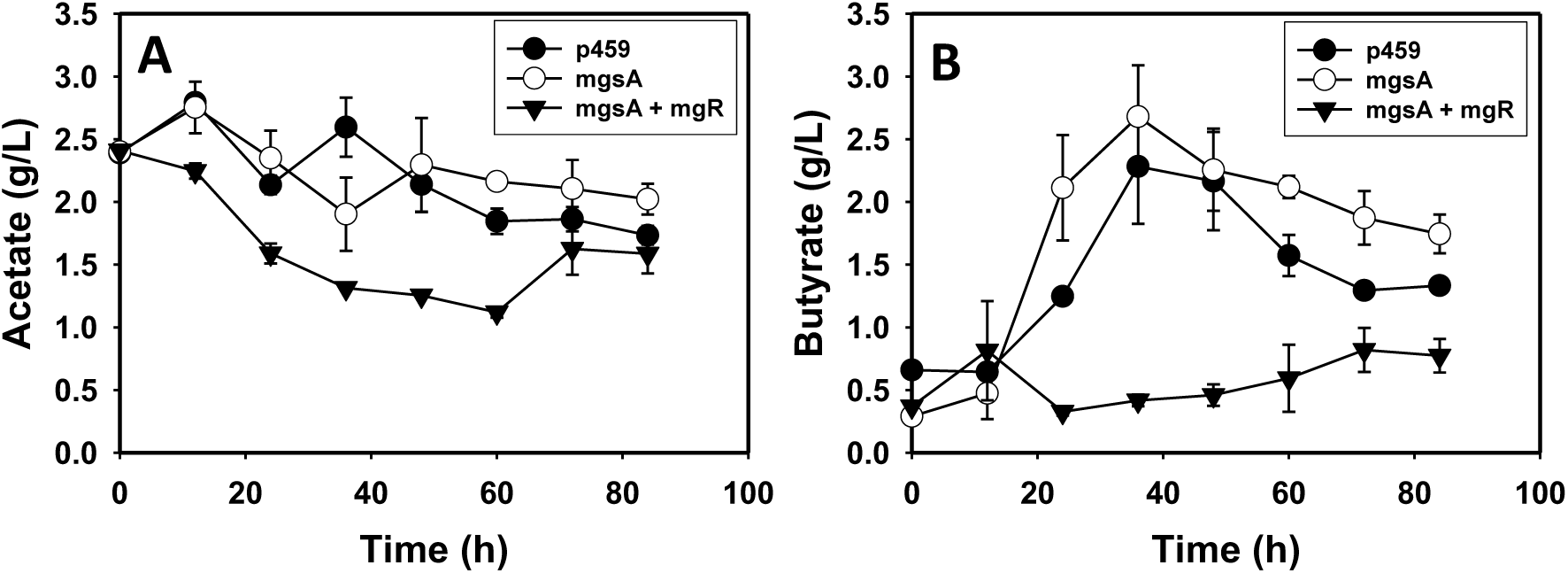
Acetate and butyrate concentrations in cultures of recombinant *Cbei* strains grown on glucose. A. Acetate concentrations; B. butyrate concentrations.

With ZnSO_4_.7H_2_O (0.01 g/L)-supplemented glucose, *Cbei*_*mgsA*+*mgR* achieved up to 21% higher OD_600 nm_ and produced at least 87% and 86% greater concentrations of butanol and ABE, respectively, when compared to *Cbei*_p459 and _*mgsA* (**Fig. 8**; **Table 4**). When lactose was used as carbon source, all three strains of *Cbei* showed reduced growth, butanol and ABE concentrations relative to the cultures grown on glucose. Nonetheless, *Cbei*_*mgsA*+*mgR* exhibited up to 35% greater OD_600 nm_ than *Cbei*_p459 and _*mgsA* (**Fig. 9A**). Similarly, the butanol and ABE concentrations of *Cbei*_*mgsA*+*mgR* were 87% and 74% and 85% and 69% greater than those of *Cbei*_p459 and _*mgsA*, respectively (**Fig. 9B and C**; **Table 4**). However, comparatively, all three strains showed similar acetate concentrations on lactose, relative to glucose-grown cultures (data not shown). Conversely, *Cbei*_*mgsA*+*mgR* exhibited slightly lower butyrate concentrations than *Cbei*_p459 and _*mgsA* in the latter stages of fermentation with lactose as carbon source (48 – 72 h; data not shown). With glucose, *Cbei*_*mgsA*+*mgR* produced 2.7- and 1.5-fold greater butanol yield and 1.1- and 1.5-fold higher ABE yield than *Cbei*_p459 and _*mgsA*, respectively (**Table 4**). Similarly, the butanol and ABE yields of *Cbei*_*mgsA*+*mgR* were at least 1.7- and 1.5-fold greater, respectively, than the yields calculated for cultures of *Cbei*_p459 and _*mgsA* when grown in the glucose + ZnSO_4_.7H_2_O medium. When lactose was the carbon source, the butanol and ABE yields of *Cbei*_*mgsA*+*mgR* were at least 1.7- and 1.4-fold higher, respectively, when compared to the yields obtained in cultures of *Cbei*_p459 and _*mgsA* (**Table 4**).

**Fig. 8.**
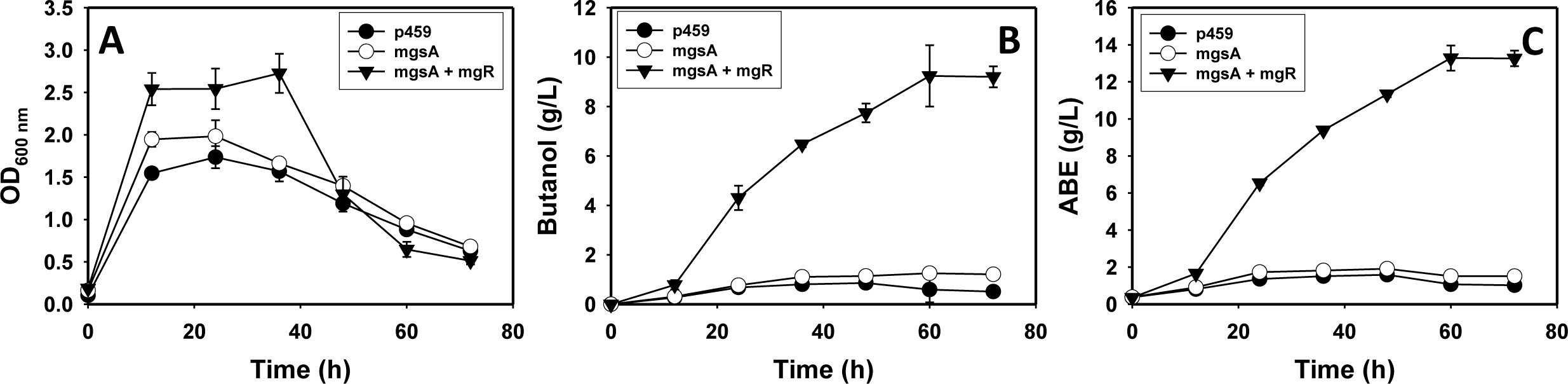
OD_600 nm_, butanol, and ABE profiles of cultures of *Cbei*_p459, _mgsA and _mgsA + mgR grown on glucose supplemented with ZnSO_4_.7H_2_O (0.01 g/L). A. OD_600 nm_; B. butanol concentrations; C. ABE concentrations.

**Fig. 9.**
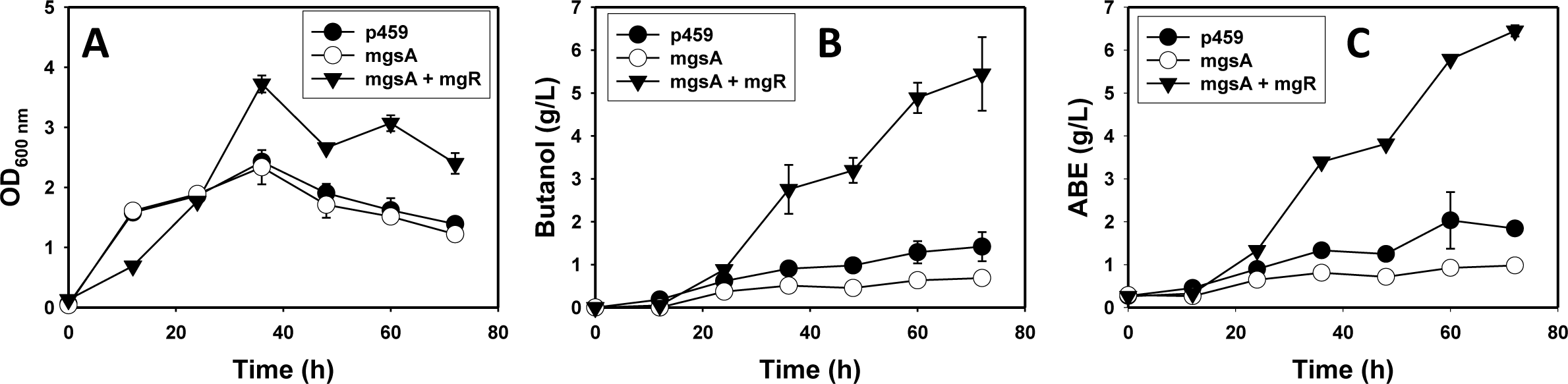
OD_600 nm_, butanol, and ABE profiles of cultures of *Cbei*_p459, _mgsA and _mgsA + mgR grown on lactose.

**Table 4.**
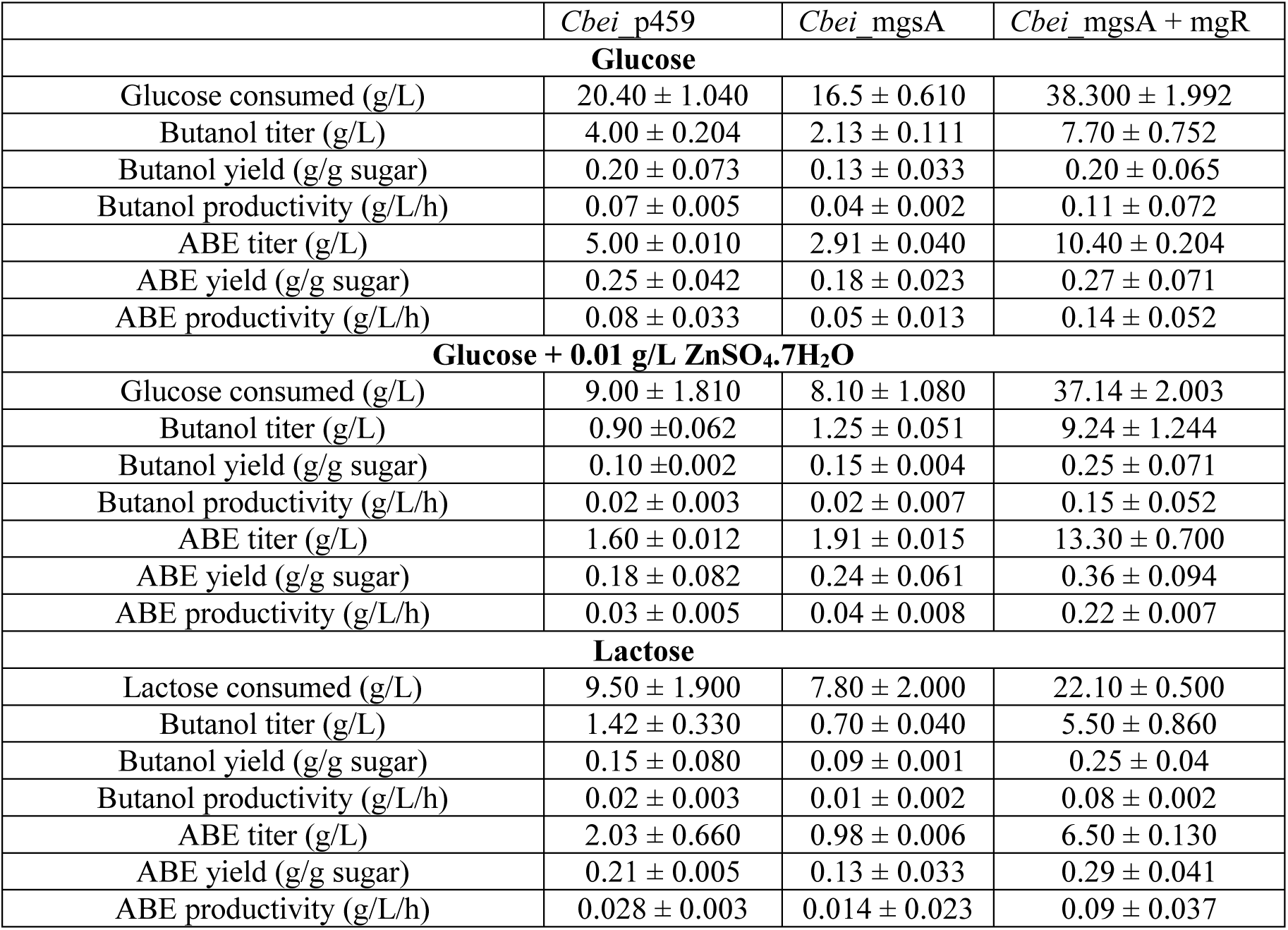
Butanol and ABE titers, yields and productivities of the recombinant strains of *C. beijerinckii* grown in different media.

|  | *Cbei_p459* | *Cbei_mgsA* | *Cbei_mgsA + mgR* |
| --- | --- | --- | --- |
| **Glucose** |  |  |  |
| Glucose consumed (g/L) | 20.40 ± 1.040 | 16.5 ± 0.610 | 38.300 ± 1.992 |
| Butanol titer (g/L) | 4.00 ± 0.204 | 2.13 ± 0.111 | 7.70 ± 0.752 |
| Butanol yield (g/g sugar) | 0.20 ± 0.073 | 0.13 ± 0.033 | 0.20 ± 0.065 |
| Butanol productivity (g/L/h) | 0.07 ± 0.005 | 0.04 ± 0.002 | 0.11 ± 0.072 |
| ABE titer (g/L) | 5.00 ± 0.010 | 2.91 ± 0.040 | 10.40 ± 0.204 |
| ABE yield (g/g sugar) | 0.25 ± 0.042 | 0.18 ± 0.023 | 0.27 ± 0.071 |
| ABE productivity (g/L/h) | 0.08 ± 0.033 | 0.05 ± 0.013 | 0.14 ± 0.052 |
| **Glucose + 0.01 g/L ZnSO<sub>4</sub>·7H<sub>2</sub>O** |  |  |  |
| Glucose consumed (g/L) | 9.00 ± 1.810 | 8.10 ± 1.080 | 37.14 ± 2.003 |
| Butanol titer (g/L) | 0.90 ± 0.062 | 1.25 ± 0.051 | 9.24 ± 1.244 |
| Butanol yield (g/g sugar) | 0.10 ± 0.002 | 0.15 ± 0.004 | 0.25 ± 0.071 |
| Butanol productivity (g/L/h) | 0.02 ± 0.003 | 0.02 ± 0.007 | 0.15 ± 0.052 |
| ABE titer (g/L) | 1.60 ± 0.012 | 1.91 ± 0.015 | 13.30 ± 0.700 |
| ABE yield (g/g sugar) | 0.18 ± 0.082 | 0.24 ± 0.061 | 0.36 ± 0.094 |
| ABE productivity (g/L/h) | 0.03 ± 0.005 | 0.04 ± 0.008 | 0.22 ± 0.007 |
| **Lactose** |  |  |  |
| Lactose consumed (g/L) | 9.50 ± 1.900 | 7.80 ± 2.000 | 22.10 ± 0.500 |
| Butanol titer (g/L) | 1.42 ± 0.330 | 0.70 ± 0.040 | 5.50 ± 0.860 |
| Butanol yield (g/g sugar) | 0.15 ± 0.080 | 0.09 ± 0.001 | 0.25 ± 0.04 |
| Butanol productivity (g/L/h) | 0.02 ± 0.003 | 0.01 ± 0.002 | 0.08 ± 0.002 |
| ABE titer (g/L) | 2.03 ± 0.660 | 0.98 ± 0.006 | 6.50 ± 0.130 |
| ABE yield (g/g sugar) | 0.21 ± 0.005 | 0.13 ± 0.033 | 0.29 ± 0.041 |
| ABE productivity (g/L/h) | 0.028 ± 0.003 | 0.014 ± 0.023 | 0.09 ± 0.037 |

## Discussion

Bioproduction of 1,2-PD has considerable environmental and economic benefits given the challenges associated with the synthetic process. However, developing an economically viable biological route for 1,2-PD production remains elusive. Because of their innate ability to produce solvents, solventogenic *Clostridium* species hold substantial promise for 1,2-PD production. With a view to establishing the capability of 1,2-PD production by the fermentative workhorse *Cbei*, we engineered two strains of this organism via heterologous and homologous expression of 1,2-PD biosynthesis genes. According to Huang et al. [26], cloning and expressing *mgsA* from *Clostridium acetobutylicum* alongside the native *gdlA* in *Escherichia coli* led to 1,2-PD production, albeit at a low concentration (3.9 mM). Similarly, cloning and expression of native *mgsA*, *gldA* and *yqhD* (aldehyde oxidoreductase) in *E. coli*, in tandem with genetic manipulation of the glycerol catabolic pathway led to the production of 5.6 g/L 1,2-PD on glycerol [27]. Notably, cloning and expression of the native *mgsA* in *Cbei* did not engender the same effect (i.e., 1,2-PD biosynthesis). Similarly, cloning and expression of *mgsA* and *mgR* from *Cpas* failed to elicit 1,2-PD biosynthesis in *Cbei*. The reason(s) for these results are not clear. However, we speculate that a stringent regulatory mechanism(s) likely limits the activity of MgsA in *Cbei*. According to Dickmanns et al. (28), in *Bacillus subtilis*, the phosphoprotein Crh exists in an unphosphorylated form in the absence of preferred carbon sources, which allows it to control the activity of MgsA by binding to the enzyme. We have not studied possible interactions between Crh and MgsA in *Cbei*. However, a Blastp search of Crh against the *Cbei* genome reveals a regulatory phosphoprotein (HPr; WP_011968555.1) involved in posttranslational modification. HPr has 43.9% similarity with Crh (with 43.9% shared identity, 96% query coverage and a score of 77.8). Indeed, investigating possible structural interactions between HPr and MgsA might help to understand the inability to *Cbei*_*mgsA* and _*mgsA*+*mgR* to produce 1,2-PD.

Given its toxicity, methylglyoxal production in *Cbei*_*mgsA* should exert a degree of stress, evidenced by impaired growth in this strain relative to the other recombinant stains studied. However, this was not the case. The growth profiles of *Cbei*_*mgsA* without methylglyoxal supplementation (**Figs. 6A, 8A & 9A**) do not suggest the presence of an added metabolic burden such as methylglyoxal production in this strain, relative to the control strain (*Cbei*_p459). This lends further weight to the notion that the recombinant MgsA in *Cbei*_*mgsA* is possibly inactive. Further, when *Cbei*_p459, _*mgsA* and _*mgsA*+*mgR* were challenged with 0.5 g/L methylglyoxal (**Fig. 2B**), *Cbei*_*mgsA* grew similarly to *Cbei*_p459 and achieved 34% higher maximum OD_600 nm_ than Cbei_*mgsA*+*mgR*.

Clearly, *Cbei*, *Cpas* and *Ctyro* possess robust innate capacities to convert acetol to 1,2-PD—a reaction catalyzed by GldA (**Fig. 1**). However, methylglyoxal is less efficiently converted to 1,2-PD in all three species. This indicates that the enzyme repertoire fundamental to methylglyoxal transformation, both via acetol and lactaldehyde are either lacking or perform extremely poorly in these organisms. Additionally, it is plausible that GldA likely does not catalyze the conversion of methylglyoxal to lactaldehyde (**Fig. 1**) in these organisms. Given the high efficiency of acetol-to-1,2-PD conversion (**Fig. 4**), if the same enzyme (i.e., GldA) catalyzes the conversion of methylglyoxal to lactaldehyde in *Cbei*, *Cpas* or *Ctyro*, either lactaldehyde and/or higer amounts of 1,2-PD might have been detected in the methylglyoxal-challenged cultures. Notably, (*S*)-(+)-1,2-PD was detected at a very low concentration in methylglyoxal-treated cultures of *Cbei* only. In fact, with 0.85 g/L methylglyoxal supplementation, only a meager 9.4% was converted to 1,2-PD (**Table 3**). While this suggests that the lactaldehyde route might be feebly active in *Cbei*, apparently, stronger competing mechanisms for methylglyoxal removal override the 1,2-PD route in this organism. It is important to highlight that whereas (*S*)-(+)-1,2-PD was detected in methylglyoxal-challenged cultures of wildtype *Cbei*, the recombinant strains did not produce (*S*)-(+)-1,2-PD. Considering the low yield of (*S*)-(+)-1,2-PD in the wildtype, the added erythromycin-mediated stress and supplementation of a lower concentration (0.5 g/L as opposed to 0.85 g/L) in cultures of the recombinant strains may have exerted further limitations on 1,2-PD biosynthesis in the recombinant strains.

Both acetol and lactaldehyde are reduced to 1,2-PD by aldehyde reductases. Our results suggest—especially with methylglyoxal challenge—that aldehyde reductases in *Cbei*, *Caps* and *Ctyro* likely have poor specificity for methylglyoxal as a substrate. Therefore, in light of the high efficiency of acetol conversion to 1,2-PD, identifying superior aldehyde reductases to reduce methylglyoxal to acetol is crucial to successful biosynthesis of 1,2-PD in *Cbei*, in particular. This alone may not prove particularly successful. This is because, in nearly all cases, the supplemented methylglyoxal was completely removed from the culture. Because of its toxicity, it is well established that the glyoxalase pathway plays a central role in the detoxification of methylglyoxal in several bacteria [29]. Therefore, the most plausible culprit for methylglyoxal detoxification in *Cbei*, *Cpas* and *Ctyro* is the glyoxalase system. The degree of methylglyoxal removal from the supplemented cultures suggest a high glyoxalase efficiency in these organism. In fact, a search of the *Cbei*, *Cpas* and *Ctyro* genomes turned up 9, 20 and 2 glyoxalases, respectively. This high level of glyoxalase redundancy, particularly in *Cbei* and *Cpas* point to a possible important role of these enzymes in both organisms. Because of their strict anaerobic nature [30-31], *Clostridium* species are particularly susceptible to oxidative damage, which might explain the preponderance of glyoxalases (in *Cbei* and *Cpas*), which play an important role in countering oxidative stress [32-33]. In the glyoxalase system, methylglyoxal is converted to *D*-lactate via *S*-*D*-lactoylgluthatione and eventually to pyruvate [29, 34]. In this study, lactate was detected in methylglyoxal-challenged cultures of *Ctyro* only, 6 h after methylglyoxal treatment (**Table 3**). It is important to note that residual methylglyoxal was only detected in cultures of *Ctyro* 6 h post supplementation. This likely indicates superior methylglyoxal detoxification machineries in *Cbei* and *Cpas*. At the time of sampling, it is possible that conversion of methylglyoxal to pyruvate may have been completed in *Cbei* and *Cpas*, both of which exhibit a high level of glyoxalase redundancy.

It is important to highlight that acetol was detected in the cultures of all the recombinant strains of *Cbei* (grown in different media), wildtype *Cbei*, Cpas and *Cytro*, without the presence of 1,2-PD, except for methylglyoxal-challenged cultures of wildtype *Cbei* (**Tables 2 & 3**). In all cases, acetol was only detectable by gas chromatography mass spectrometry but not by gas chromatography alone. We suspect that only traces of acetol may have been produced in these cultures, which were possibly too low to elicit further conversion to 1,2-PD. When we added 0.2 g/L acetol to cultures of *Cbei*, *Cpas* and *Ctyro*, it was completely reduced to 1,2-PD, which supports the assertion that much lower concentrations were likely produced in the cultures listed above.

The butanol and ABE concentrations of *Cbei*_*mgsA*+*mgR* in all conditions tested implicate the MgR from *Cpas* in butanol biosynthesis (**Figs. 6, 8 & 9**). Further, since we did not challenge *Cbei*_*mgsA*+*mgR* with glyoxal, we cannot rule out the ability of the enzyme product of this gene to reduce glyoxal. However, inability of *Cbei*_*mgsA*+*mgR* to produce 1,2-PD in methylglyoxal-supplemented cultures coupled with the removal of methylglyoxal from the cultures, led us to conclude that the enzyme encoded by *Cpas mgR* does not utilize methylglyoxal as a substrate. Even in the presence of high levels of glyoxalase expression and activity, kinetically, this should not completely eliminate methylglyoxal conversion to acetol and then to 1,2-PD in *Cbei*_*mgsA*+*mgR*. More so, the high efficiency of acetol conversion to 1,2-PD by both the recombinant *Cbei* strains and the wildtype (**Tables 2 & 3; Fig. 4**) highlight the inability of *Cpas* MgR to catalyze methylglyoxal reduction to acetol. Furthermore, detection of (*S*)-(+)-1,2-PD in the cultures of wildtype *Cbei* treated with methylglyoxal suggests that *Cpas* MgR does not convert methylglyoxal to lactaldehyde either (**Table 3**). Both *mgsA* and *mgR* were cloned under the control of *Cbei* P*adc*. P*adc* is induced in the early solventogenic phase (10 – 12 h), characterized by the onset of solvent (ABE) biosynthesis. In all cases, acetone, butanol and ethanol were detected in the cultures of *Cbei*_*mgsA*+*mgR* at 12 h, when methylglyoxal supplementation took place. Obviously, expression of *mgR* should be active at this time point, which is evidenced by 34% and 190% higher butanol titers at 12 h in cultures of *Cbei*_*mgsA*+*mgR* relative to those of *Cbei*_p459 grown on glucose and zinc sulfate-supplemented glucose, respectively (**Figs. 6 & 8**; methylglyoxal was supplemented in glucose-grown cultures). *Cbei*_*mgsA* exhibited even greater delay in solvent biosynthesis. Evidently, the MgR from *Cpas* does not appear to act on methylglyoxal. On the other hand, with either glucose or lactose, *Cbei*_*mgsA*+*mgR* produced significantly more butanol and ABE than *Cbei*_*mgsA* and *Cbei*_p459. Additionally, it grew better and exhibited superior acid re-assimilation—especially on glucose—relative to *Cbei*_*mgsA* and *Cbei*_p459. The MgR from *Cpas* is annotated as an aldo-keto reductase (AKR). Generally, AKRs reduce ketones and aldehydes to alcohol and tend to exhibit broad substrate ranges with substrates including glucose, glycosylation products and pollutants [35].

With multiple substrates as candidates between glucose and butanol in the ABE pathway, it is not clear where the *Cpas* MgR exerts its effect leading to increased butanol and ABE production in *Cbei*_*mgsA*+*mgR*. However, it is obvious that the recombinant MgR in *Cbei*_*mgsA*+*mgR* acts on a central substrate(s) in this pathway, likely expediting flux towards butanol. Cloning, expression, purification and characterization of the *Cpas* MgR will shed more light on how it mediates increased butanol and ABE biosynthesis in *Cbei*_*mgsA*+*mgR*. Previously, we have shown that chromosomal integration and expression of an AKR (encoded by the open reading frame Cbei_3974) in *Cbei* led to enhanced furfural detoxification and ABE production [21]. More importantly, the strain expressing the AKR produced more butanol even in cultures un-supplemented with furfural. It appears some AKRs likely act on intermediates such a glyceraldehyde-3-phosphate, acetaldehyde or butyraldehyde in the ABE pathway. Notably, increase in butanol production was more significant than the increase observed for ethanol. In addition, enhanced butanol biosynthesis and acid re-assimilation tend to occur in tandem in solventogenic *Clostridium* species [36]. Thus, if the *Cpas* MgR utilizes acetaldehyde as substrate, significantly higher increases in ethanol should have been observed, which was not the case. On the other hand, given the natural tendency of solventogenic clostridia to produce more butanol, increased utilization of glyceraldehyde-3-phosphate might still favor butanol biosynthesis. However, this does not explain greater assimilation of acids in *Cbei*_*mgsA*+*mgR* (**Fig. 7**). Acid re-assimilation is more directly tethered to butanol biosynthesis, which suggests that butyraldehyde might be a plausible substrate for the MgR from *Cpas*. Taken together, the MgR from *Cpas* represents a promising candidate for increasing butanol biosynthesis in solventogenic clostridia. It is important to highlight that all the constructs described here are plasmid-borne, which required erythromycin supplementation to maintain the plasmids. Ultimately, this limits growth and ABE production. Therefore, most likely, chromosomal integration of *mgR* might further increase butanol and ABE yield.

## Conclusions

In this study, we engineered strains of *Cbei* for 1,2-PD production and investigated the bottlenecks to 1,2-PD biosynthesis in this organism. Furthermore we investigated the metabolic impediments to 1,2-PD production in *Cpas* and *Ctyro*. Our results show that vastly inefficient reduction of methylglyoxal severely impairs 1,2-PD production in these organisms. Additionally, cloning and expression of *Cpas mgR* in *Cbei* led to the production of up to 88% higher butanol concentration in the resulting strain, relative to the control and the strain of *Cbei* expressing the native *mgsA*.

## Ethics approval and consent to participate

Not applicable

## Consent for publication

I, Victor Ujor do hereby, give my consent for the publication of identifiable details of this manuscript, including the figures within the text to be published in Biotechnology for Biofuels and Bioproducts.

## Availability of data and materials

The datasets supporting the conclusions of this article are included within the article.

## Competing interests

The authors declare that they have no competing interests.

## Funding

This work was supported by funding from USDA-National Institute of Food and Agriculture (Hatch award, grant no. WIS04018) and the Dairy Innovation Hub (AAK5475) to VCU.

## Authors’ contributions

EG co-designed and conducted the fermentation experiments, contributed to the construction of the recombinant strains, conducted chromatographic analyses, analyzed and interpreted the results, read and corrected the manuscript draft. SK constructed the recombinant strain, analyzed and interpreted the results, read and edited the manuscript draft. VCU conceived the study and obtained funding for the work, co-designed and facilitated the experiments, analyzed and interpreted the results and wrote the manuscript draft.

## Acknowledgements

We thank Dr. Thaddeus Ezeji (Dept. of Animal Sciences, The Ohio State University) for kindly providing us with *Clostridium beijerinckii* NCIMB 8052 and *Clostridium pasteurianum* ATCC 6013 and Dr. Yi Wang (Biosystems Engineering, Auburn University) for kindly supplying the *Clostridium tyrobutyricum* ATCC 25755 used in this study.

## Notes

### Competing Interest Statement

The authors have declared no competing interest.

